# Autophagosome turnover requires Arp2/3 complex-mediated maintenance of lysosomal integrity

**DOI:** 10.1101/2024.03.12.584718

**Authors:** Corey J. Theodore, Lianna H. Wagner, Kenneth G. Campellone

## Abstract

Autophagy is an intracellular degradation process that maintains homeostasis, responds to stress, and plays key roles in the prevention of aging and disease. Autophagosome biogenesis, vesicle rocketing, and autolysosome tubulation are controlled by multiple actin nucleation factors, but the impact of actin assembly on completion of the autophagic pathway is not well understood. Here we studied autophagosome and lysosome remodeling in fibroblasts harboring an inducible knockout (iKO) of the Arp2/3 complex, an essential actin nucleator. Arp2/3 complex ablation resulted in increased basal levels of autophagy receptors and lipidated membrane proteins from the LC3 and GABARAP families. Under both steady-state and starvation conditions, Arp2/3 iKO cells accumulated abnormally high numbers of autolysosomes, suggesting a defect in autophagic flux. The inability of Arp2/3 complex-deficient cells to complete autolysosome degradation and turnover is explained by the presence of damaged, leaky lysosomes. In cells treated with an acute lysosomal membrane-damaging agent, the Arp2/3-activating protein WHAMM is recruited to lysosomes, where Arp2/3 complex-dependent actin assembly is crucial for restoring intact lysosomal structure. These results establish the Arp2/3 complex as a central player late in the canonical autophagy pathway and reveal a new role for the actin nucleation machinery in maintaining lysosomal integrity.

## INTRODUCTION

Autophagy is a fundamental intracellular catabolic mechanism in which cytoplasmic material is captured by double membrane-bound sequestering vesicles called autophagosomes that fuse with lysosomes for degradation and recycling (Ohsumi, 2014). This process maintains homeostasis and allows cells to respond to multiple stressors (Morishita and Mizushima, 2019). In a canonical pathway of autophagy, nutrient starvation results in activation of a signaling cascade that leads to accumulation of the phospholipid PI(3)P at ER-associated omegasomes, where autophagosome biogenesis takes place (Palamiuc et al., 2020; Roberts and Ktistakis, 2013). Formation of the autophagosomal isolation membrane, termed a phagophore, involves lipidation of the LC3 and GABARAP subclasses of ATG8-family proteins and their incorporation into the double membrane (Nguyen and Lazarou, 2022). In selective forms of autophagy, ATG8 proteins are linked to ubiquitinated protein targets through autophagy cargo receptors, which contain LC3-interacting regions and ubiquitin-binding domains (Johansen and Lamark, 2020). Following membrane closure, mature autophagosomes undergo fusion with lysosomes, and both the inner autophagosomal membrane and autophagic cargo are degraded (Nakamura and Yoshimori, 2017). To regenerate lysosomes, autolysosome-derived membrane tubules take part in a process called autophagic lysosome reformation (Yim and Mizushima, 2020).

The actin cytoskeleton impacts practically every cellular function (Campellone et al., 2023; Kramer et al., 2022), and its participation in autophagy was initially found to be important for autophagosome formation during starvation (Aplin et al., 1992). The Arp2/3 complex, an essential actin nucleator, was later shown to be crucial for selective autophagy in yeast (Monastyrska et al., 2008). The role of actin assembly in autophagy has since become best characterized during autophagosome biogenesis in mammalian cells. For instance, the Arp2/3 complex localizes to developing phagophores, and the actin filament capping protein CapZ controls phagophore size and shape in a PI(3)P-dependent manner (Mi et al., 2015).

Throughout the canonical autophagy pathway, the Arp2/3 complex is activated by actin nucleation-promoting factors from the Wiskott-Aldrich Syndrome Protein (WASP) family. As examples, the WASP-family proteins WHAMM and JMY localize to omegasomes, phagophores, and autophagosomes, and Arp2/3 activation by each protein contributes to autophagosome biogenesis and maturation (Coutts and La Thangue, 2015; Kast et al., 2015). WHAMM is recruited to PI(3)P early in the autophagy pathway (Mathiowetz et al., 2017), while JMY interacts with LC3B, a member of the ATG8 family (Hu and Mullins, 2019). WHAMM is also important for linking cargo receptors to ATG8-family proteins during autophagosome closure (Coulter et al., 2024). Once autophagosomes are formed, WHAMM and JMY are both able to stimulate the Arp2/3 complex to drive actin-based rocketing (Hu and Mullins, 2019; Kast et al., 2015). Another WASP-family member, WASH, mediates endosomal membrane trafficking and can have either stimulatory or inhibitory effects on autophagosome formation (Xia et al., 2013; Zavodszky et al., 2014).

Actin assembly additionally influences later stages of the autophagic pathway. The atypical Arp2/3 activator Cortactin controls actin branching during autophagosome-lysosome fusion (Lee et al., 2010). WHAMM contains two PI(4,5)P2-binding motifs that allow localization to autolysosomes, where it promotes actin assembly to support lysosomal membrane tubulation (Dai et al., 2019). WHAMM also interacts with a member of the BLOC-1 lysosome biogenesis complex (Wu et al., 2021). WASH can associate with the BLOC-1 complex as well (Monfregola et al., 2010; Ryder et al., 2013), and WASH depletion has harmful effects on lysosomal populations (Gomez et al., 2012). Thus, numerous actin nucleation-promoting factors direct the Arp2/3 complex to polymerize actin at numerous steps in the autophagic pathway, from phagophore biogenesis through autophagic lysosome reformation.

Due to this complicated nature of cytoskeletal regulation by multiple nucleation-promoting factors, the variability in autophagy phenotypes depending on cell type, and the different approaches for manipulating Arp2/3 functions, the overall consequence of Arp2/3 complex inactivation on autophagy has been difficult to establish. To determine the net outcome of removing the Arp2/3 complex completely, we utilized a conditional Arp2/3 complex knockout cell model (Haarer et al., 2023; Rotty et al., 2015). Our findings reveal that the Arp2/3 complex is most important late in autophagy, because Arp2/3-mediated actin assembly maintains lysosome integrity and enables lysosomal membrane repair.

## RESULTS

### The Arp2/3 complex is required for autophagic membrane turnover

Multiple nucleation-promoting factors including WHAMM, JMY, WASH, and Cortactin participate in assembling actin filaments at phagophores, autophagosomes, or autolysosomes (Campellone et al., 2023; Kramer et al., 2022). These observations led us to question where and when the Arp2/3 complex has the greatest impact on autophagy. To address this problem while avoiding the confounding effects of partial loss-of-function or isoform-specific inhibition, we studied autophagy using inducible ArpC2 knockout fibroblasts (Rotty et al., 2015). These cells contain a floxed *Arpc2* allele and encode the CreER recombinase. The ArpC2 subunit is essential for forming a functional Arp2/3 complex, so exposure to 4-hydroxytamoxifen (4-OHT) causes the permanent loss of ArpC2 and leads to depletion of the entire complex with kinetics that have been characterized previously (Haarer et al., 2023). In our current study, cells carrying the conditional *Arpc2* allele were treated with DMSO to maintain a control (Flox) cell population or with 4-OHT to generate induced knockout (iKO) cells, and experiments were performed 6-9 days after initiation of treatment, when iKO cells are completely devoid of the Arp2/3 complex.

Mammalian cells encode several selective autophagy receptors whose expression levels are inversely related to the amount of autophagic degradation or flux. To determine the effects of Arp2/3 complex deletion on autophagy receptor levels, we immunoblotted Flox and iKO cell extracts with antibodies to the prominent autophagy receptors NBR1, TAX1BP1, Optineurin, SQSTM1/p62, and NDP52. Compared to the tubulin, actin, and GAPDH loading controls, the levels of NBR1, TAX1BP1, Optineurin, and p62 were noticeably higher in iKO cells (Figure 1A). Quantification revealed that Optineurin and p62 were 2-3-fold more abundant in iKO cells than in Flox cells, while NDP52 quantities were similar between the two cell types (Figure 1B). These results imply that autophagic flux is inefficient in Arp2/3 complex iKO cells.

**Figure 1.**
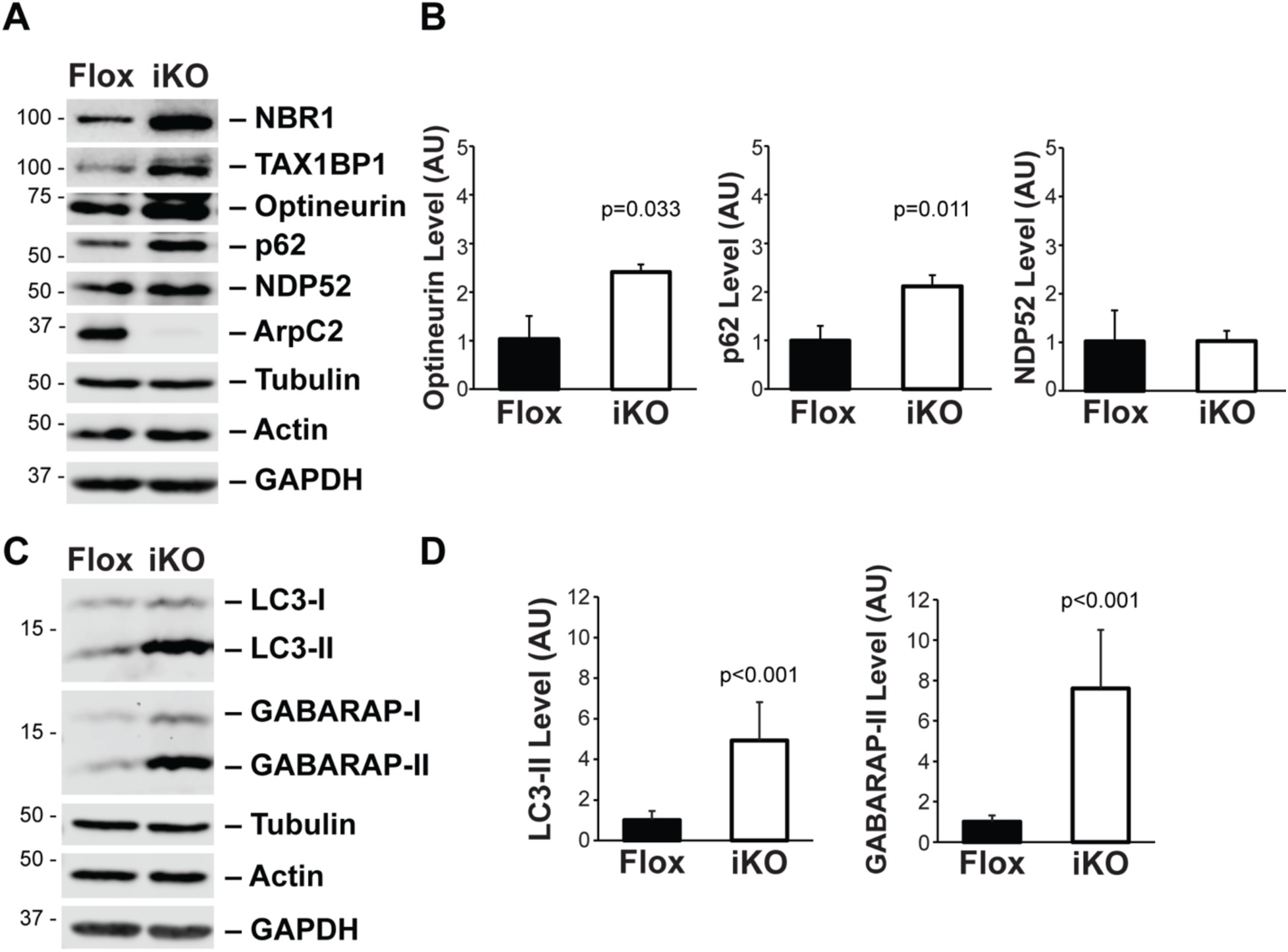
Autophagy receptors and lipidated ATG8-family proteins accumulate in Arp2/3 complex induced knockout (iKO) cells. **(A)** Mouse tail fibroblasts (MTFs) were treated with DMSO (Flox) or 4-OHT (iKO) for 6d and collected at 8d. Samples were then lysed, subjected to SDS-PAGE, and immunoblotted with antibodies to NBR1, TAX1BP1, Optineurin, SQSTM1/p62, NDP52, ArpC2, Tubulin, Actin, and GAPDH. **(B)** For quantification, the relative amounts of Optineurin, p62, and NDP52 were calculated by dividing their respective band intensities by the average loading control intensities. Each bar represents the mean ±SD from n = 3-5 experiments. AU, arbitrary units. **(C)** Flox and iKO cell extracts were collected at 8d and immunoblotted with antibodies to LC3, GABARAP, and the loading controls. **(D)** For quantification, the relative amounts of LC3-II and GABARAP-II were calculated by dividing their respective band intensities by the average loading control intensities. Each bar represents the mean ±SD from 3-5 experiments.

Autophagy receptors are linked to autophagic membranes through interactions with ATG8-family proteins. In mammals, the ATG8 family includes three LC3s (LC3A/B/C) and three GABARAPs (GABARAP, GABARAP-L1/L2), which exist as immature forms (e.g., LC3-I, GABARAP-I) that are cytosolic, and mature phosphatidylethanolamine-conjugated forms (e.g., LC3-II, GABARAP-II) that are integral to autophagosomal membranes (Klionsky et al., 2021; Mizushima, 2020). To next determine the effects of Arp2/3 complex deletion on ATG8 abundance and lipidation, we immunoblotted Flox and iKO cell extracts with polyclonal antibodies that recognize the LC3 and GABARAP protein subclasses. The levels of both LC3-II and GABARAP-II were clearly higher in iKO cells than in Flox cells (Figure 1C). Quantification indicated that the amounts of lipidated LC3 and GABARAP were, on average, 5-8-fold higher in iKO cells (Figure 1D). These data suggest that autophagic membrane protein turnover is impaired in the absence of the Arp2/3 complex.

LC3-II and GABARAP-II levels are considered to be indicative of the amount of autophagosomal membranes in cells (Klionsky et al., 2021; Mizushima, 2020). To directly visualize autophagosome abundance, we fixed the Flox and iKO cells, stained them with antibodies to LC3 or GABARAP, and examined them by fluorescence microscopy. Compared to Flox cells, in which LC3 and GABARAP staining was diffuse or occasionally punctate, iKO cells more frequently possessed larger, ring-like LC3 and GABARAP structures (Figure 2A and 2B). These observations show that Arp2/3-deficient cells are more likely to accumulate mature autophagosomal organelles.

**Figure 2.**
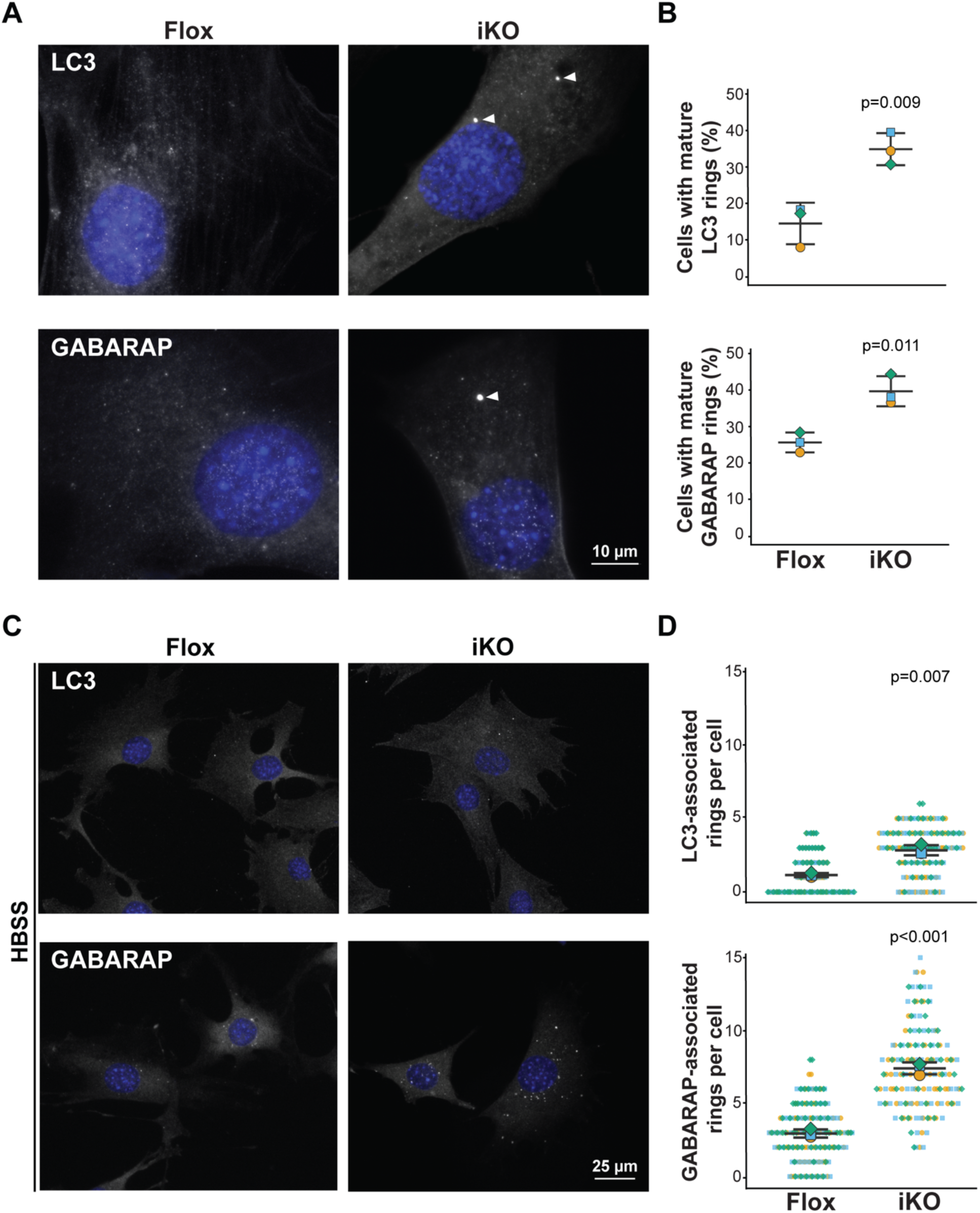
Deletion of the Arp2/3 complex leads to an increase in mature autophagic rings in fed and starved cells. **(A)** MTFs were treated with DMSO (Flox) or 4-OHT (iKO) for 6d and fixed at 8d. Cells were then stained with antibodies to detect LC3 or GABARAP (gray) and with DAPI to visualize DNA (blue). Arrowheads highlight mature autophagosomal rings. **(B)** The percentage of cells with mature rings was quantified. Large symbols represent the average %s from individual experiments in which 42-61 cells were examined. Lines denote the mean ±SD from n = 3 experiments. **(C)** Flox and iKO cells were treated with HBSS for 4h on day 8, fixed, and stained as in part (A). **(D)** Superplots depict the number of mature autophagosomal rings per cell. Small symbols represent the #s within individual cells. Large symbols represent the average #s from individual experiments in which 45-67 cells were examined. Lines denote the mean ±SD from n = 3 experiments.

To determine if this phenotype would persist under a stress condition that induces autophagy, we starved Flox and iKO cells in Hank’s Balanced Salt Solution (HBSS) prior to performing immunofluorescence microscopy. As expected, punctate LC3 and GABARAP staining increased following starvation (Figure 2C). Moreover, compared to Flox cells, iKO cells contained about twice as many mature autophagosomal rings (Figure 2D). To corroborate these results with an independent autophagy-inducing stressor, we insulted Flox and iKO cells with Carbonyl cyanide m-chlorophenyl hydrazone (CCCP), which causes a decrease in mitochondrial membrane potential and stimulates both mitophagy and other forms of autophagy. Again, the iKO cells contained significantly more LC3- and GABARAP-associated autophagosomal rings (Figure S1). Taken together, the above findings are consistent with the conclusion that the Arp2/3 complex is crucial for autophagosomal membrane turnover.

While it was possible that enhanced autophagosome biogenesis could give rise to the numerous mature autophagosomal structures in Arp2/3 iKO cells, this seemed unlikely, because several lines of evidence indicate that the Arp2/3 complex functions in promoting autophagosome biogenesis. For example, the Arp2/3-activating proteins WHAMM and JMY have been shown to increase the kinetics of LC3 lipidation, enlargement of phagophores, and maturation and closure of LC3-associated membranes (Coulter et al., 2024; Coutts and La Thangue, 2015; Kast et al., 2015; Mathiowetz et al., 2017). Nevertheless, to be certain that the increase in autophagosomal structures in Arp2/3 complex-deficient cells was not due to more autophagosome biogenesis, we treated Flox and iKO cells with chloroquine, which inhibits lysosome acidification and blocks autophagosome degradation (Mauthe et al., 2018; Poole and Ohkuma, 1981). If biogenesis was increased in iKO cells, then chloroquine should create a greater disparity in autophagosome numbers between Flox and iKO cells, but the quantities of LC3 and GABARAP rings in chloroquine-treated Flox and iKO cells were indistinguishable from one another (Figure 3A and 3B). Consistent with an actual decrease in autophagosome formation in iKO cells, calculations of the fold-change in autophagosome numbers pre- and post-chloroquine exposure gave lower values for the iKO cells compared to the Flox cells (Figure 3C). Thus, in Arp2/3 complex knockout cells, a modest inefficiency in autophagosome biogenesis is masked by a dominant defect in autophagosome turnover.

**Figure 3.**
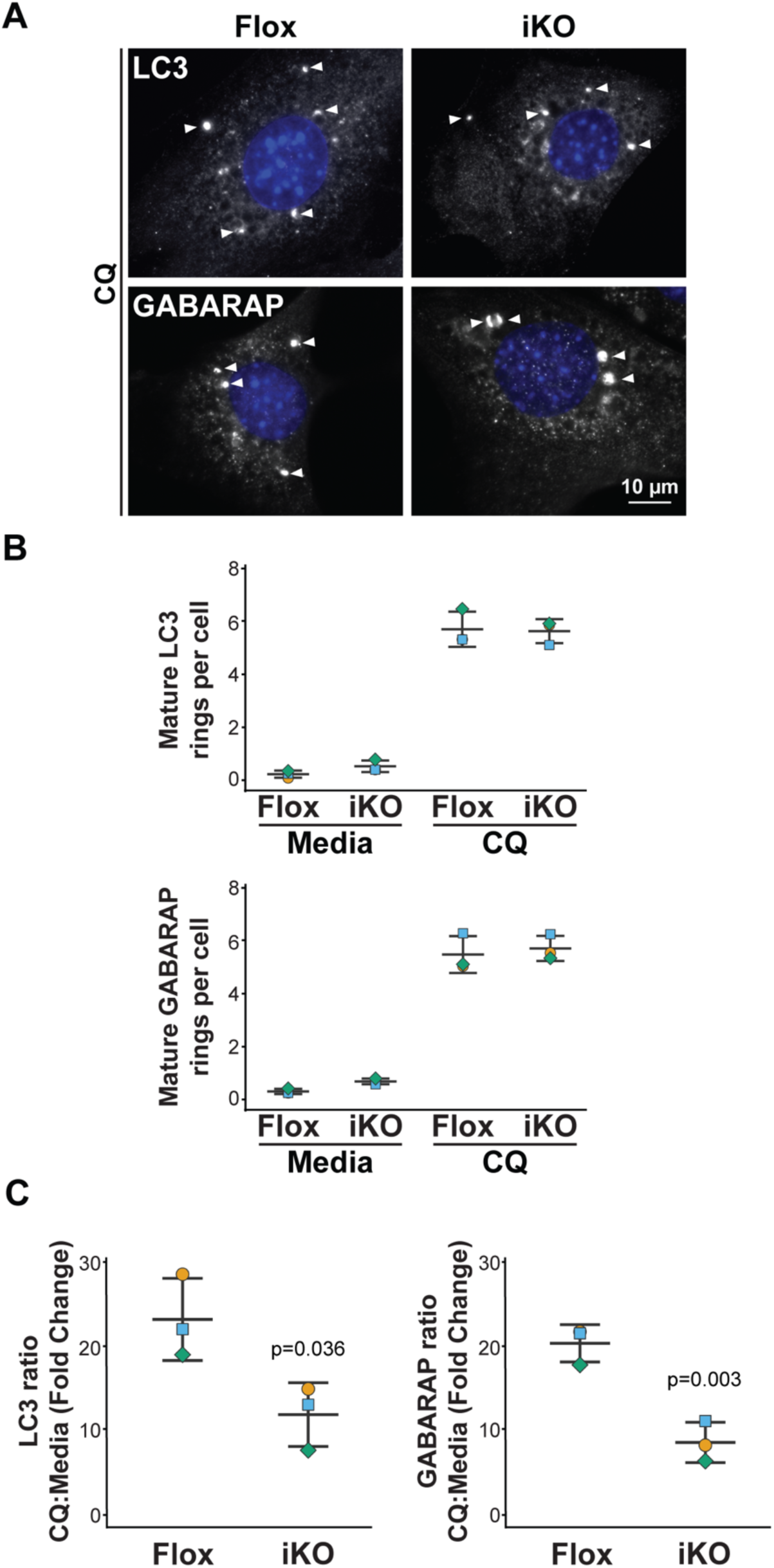
The increase in autophagic membranes in iKO cells is not due to an increase in autophagosome biogenesis. **(A)** Flox and iKO cells were treated with chloroquine (CQ) for 2h on day 8, fixed, and stained with antibodies to LC3 or GABARAP (gray) and DAPI (blue). Arrowheads highlight mature autophagosomal rings. **(B)** The number of mature rings per cell was quantified. Large symbols represent the average #s from individual experiments in which 30-42 cells were examined. Lines denote the mean ±SD from n = 3 experiments. **(C)** The fold change in the # of mature rings per cell following CQ treatment was calculated by dividing the value for CQ-treated cells by the value for control media-treated cells. Each large symbol represents the fold change in one experiment, and lines denote the mean ±SD from n = 3 experiments.

### Autophagy impairment in Arp2/3 complex iKO cells is due to lysosomal damage

To better characterize why the Arp2/3 complex-deficient cells were unable to degrade LC3- and GABARAP-associated membranes, we next sought to determine whether the accumulation of autophagosomal ring structures was caused by a defect before or after autophagosome-lysosome fusion. To do this, we transfected Flox and iKO cells with a RFP-GFP tandem fluorescent LC3 (tfLC3) construct, which allows for a visual differentiation of autophagosomes and autolysosomes (Kimura et al., 2007). Both GFP and RFP fluoresce when LC3 is associated with phagophores and autophagosomes, but once autophagosomes fuse with lysosomes to form autolysosomes, the low lysosomal pH quenches GFP fluorescence, leaving only RFP visible. Examination of transfected Flox and iKO cells under normal steady-state conditions revealed that the two cell types harbored similar numbers of autophagosomes, but that iKO cells had significantly more autolysosomes (Figure 4A and 4B). These results indicate that autophagosomes and lysosomes can associate with one another in the absence of the Arp2/3 complex, but that the autophagy pathway is stalled at the autolysosome degradation step.

**Figure 4.**
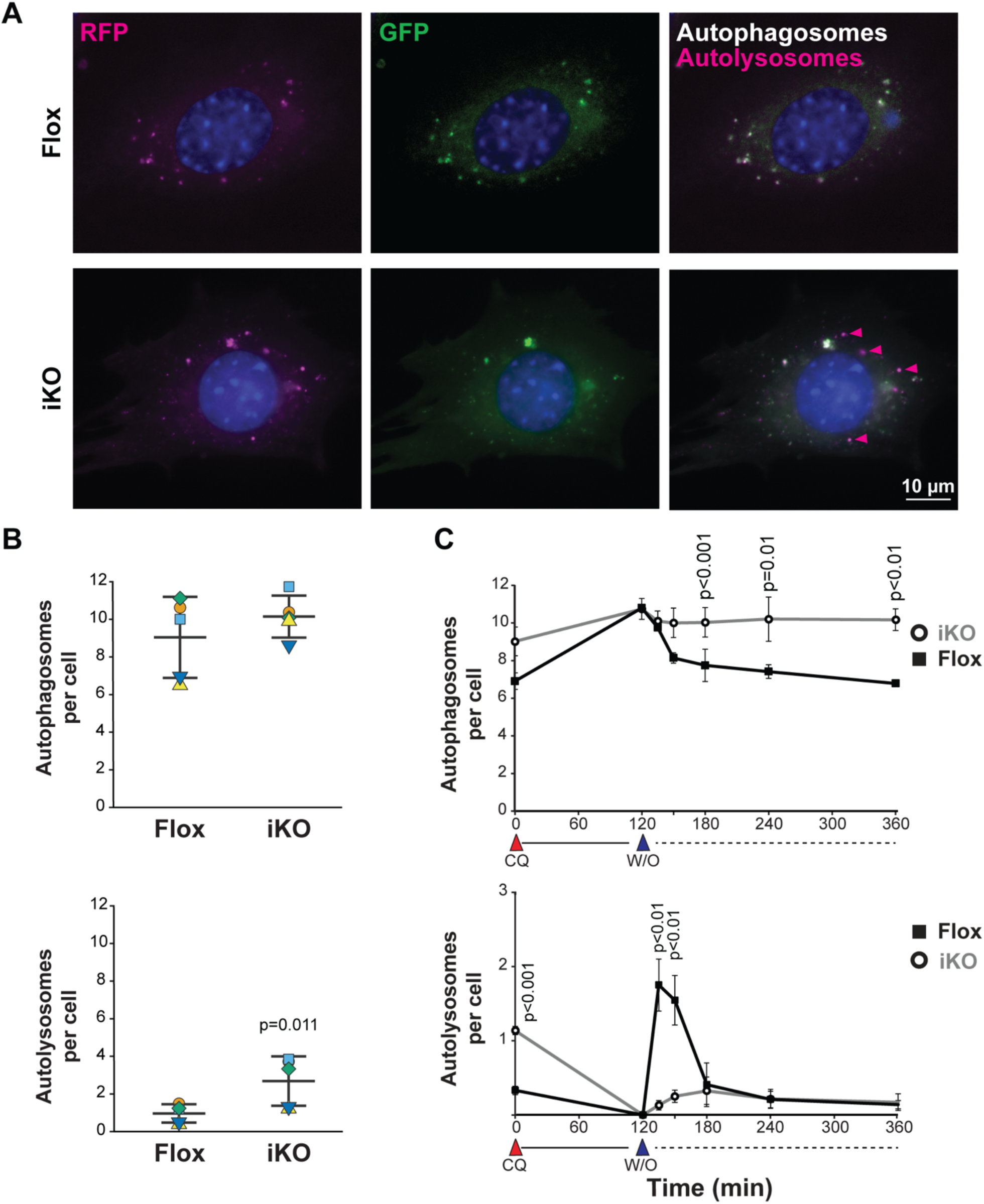
Arp2/3 complex depletion causes a defect in autolysosome turnover. **(A)** MTFs were treated with DMSO (Flox) or 4-OHT (iKO) for 2d, transfected with RFP-GFP-LC3, fixed at 6d, and stained with DAPI (blue). Autophagosomes exhibit both GFP (green) and RFP (magenta) fluorescence (white when merged), whereas autolysosomes display only RFP (magenta) fluorescence. Arrowheads highlight autolysosomes in an iKO cell. **(B)** The numbers of autophagosomes and autolysosomes per cell were quantified. Large symbols represent the average #s from individual experiments in which 10-23 cells were examined. Lines denote the mean ±SD from n = 5 experiments. **(C)** Flox and iKO cells were treated with chloroquine (CQ) for 2h on day 6, subjected to a media washout (W/O), fixed at the indicated timepoints, and imaged. Each symbol represents the mean # of autophagosomes or autolysosomes ±SD from n = 3 experiments.

To investigate the kinetics of autophagosome turnover, we treated tfLC3-expressing Flox and iKO cells with chloroquine to cause a buildup of autophagosomes before washing out the drug and monitoring the appearance and clearance of autolysosomes. After two hours of chloroquine exposure, the number of autophagosomes was high and autolysosomes were absent in both Flox and iKO cells, as expected (Figure 4C). Within 10-20 minutes after washout, the Flox cells exhibited a sharp decrease in autophagosome quantities and a corresponding spike in autolysosome numbers (Figure 4C). In the 60-120 minute washout time range, the Flox cells then returned to their steady state levels of autophagosomes and autolysosomes (Figure 4C). The iKO cells, in contrast, demonstrated a minimal dip in autophagosome quantity and only a slight rise in autolysosome numbers during a 4 hour washout period (Figure 4C). Taken together, these observations show that the increase of autophagic organelles in Arp2/3 complex-deficient cells is caused by a lysosomal defect.

Because the quenching of GFP was so poor in iKO cells after chloroquine washout, we wondered if lysosomal acidity was altered in the Arp2/3-depleted cells. Consistent with this idea, recent work using Flox and iKO cells showed that LysoTracker, a fluorescent probe which labels acidic intracellular structures, stained discrete circular lysosomes in Flox cells, but broadly stained the cytoplasm when iKO cells were senescent (Haarer et al., 2023). This pattern of intensely-focused lysosome staining in Flox cells versus widely-distributed cytoplasmic staining in iKO cells was also apparent at the earlier timepoint used in our experiments (Figure 5A).

**Figure 5.**
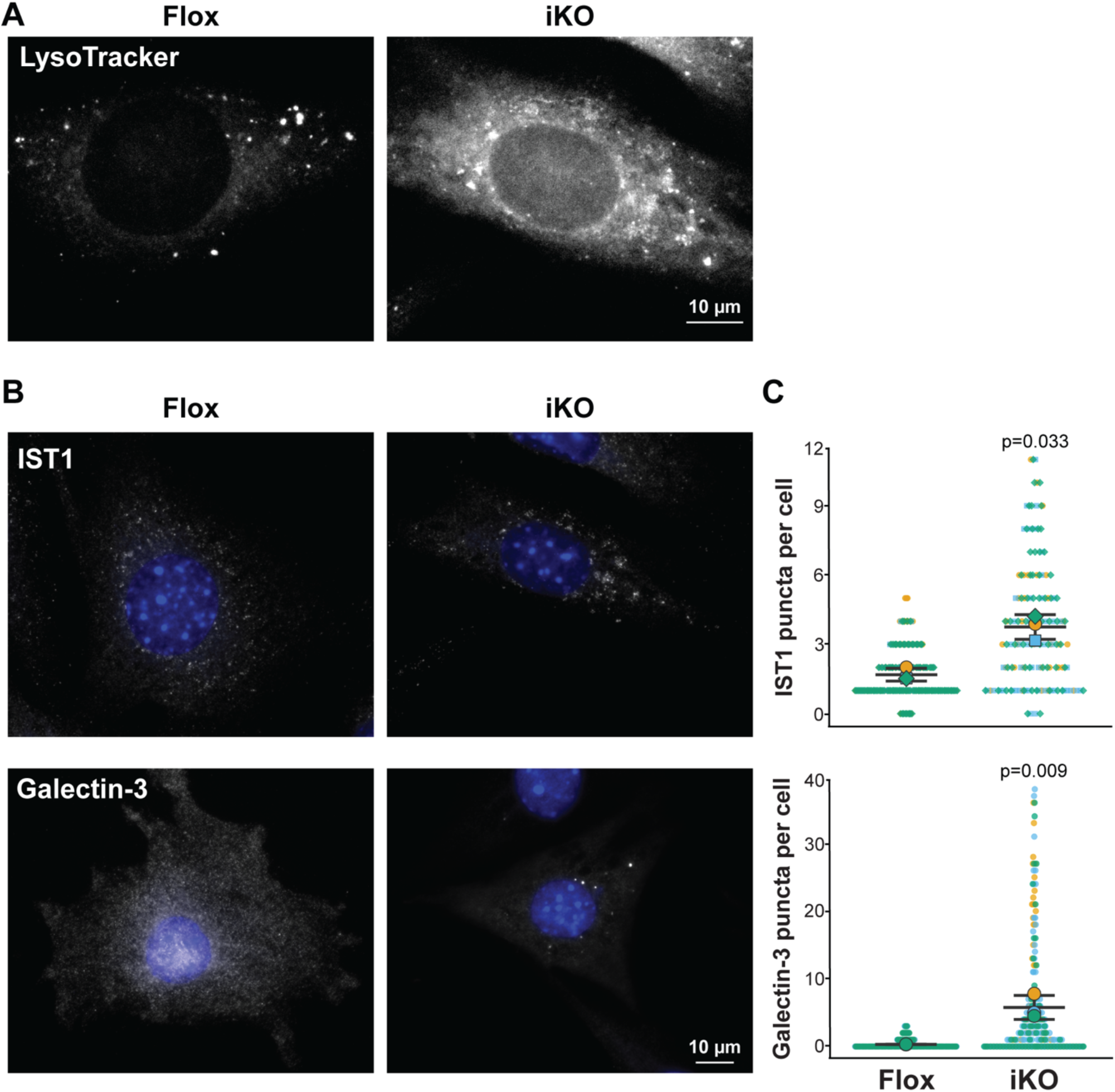
Lysosomal damage is increased when the Arp2/3 complex is deleted. **(A)** Flox and iKO cells were treated with LysoTracker for 30min and fixed at 8d. **(B)** Cells were stained with antibodies to IST1 or Galectin-3 (gray) and DAPI (blue). **(C)** The numbers of bright IST1 or Galectin-3 puncta per cell were quantified. Small symbols represent the #s within individual cells. Large symbols represent the average #s from individual experiments in which 48-85 cells were examined. Lines denote the mean ±SD from n = 3 experiments.

We hypothesized that the more diffuse LysoTracker staining in the iKO cells could be due to the presence of leaky, damaged lysosomes. To assess lysosomal membrane damage, we stained Flox and iKO cells with antibodies to IST1 and Galectin-3. IST1 is a subunit of the ESCRT-III complex that mediates repair of mildly damaged lysosomal membranes (Radulovic et al., 2018), while Galectin-3 is a glycan-binding protein that is recruited to the inner lysosomal membrane following more severe damage where it coordinates repair responses or targeting by the lysophagy machinery (Jia et al., 2020). IST1 staining was diffuse or dimly punctate in Flox cells but was brightly punctate in iKO cells (Figure 5B). Similarly, Galectin-3 localization was mostly diffuse in Flox cells but was in larger, brighter spots in iKO cells (Figure 5B). Quantification of the number of intense IST1 and Galectin-3 puncta per cell confirmed that significantly more lysosomes were damaged when the Arp2/3 complex was not present (Figure 5C). Thus, the autophagic dysfunction in Arp2/3-deficient cells is most likely due to a lack of lysosomal integrity.

### WHAMM-directed, Arp2/3 complex-mediated actin assembly supports lysosomal repair

Since the primary biochemical function of the Arp2/3 complex is to nucleate actin, we next wanted to ascertain whether actin was recruited to damaged lysosomes. To do this, we generated Flox and iKO cells stably expressing the lysosome marker LAMP1-RFP and treated them briefly (for 15 minutes) with the lysosome-damaging agent L-Leucyl L-Leucine Methyl ester (LLOMe) hydrobromide (Aits et al., 2015; Thiele and Lipsky, 1990). The cells were then fixed and stained with antibodies to Galectin-3 to identify the damaged lysosomes and with phalloidin to visualize actin filaments. In Flox cells, Galectin-3-positive RFP-LAMP1 vesicles were associated with actin, while in iKO cells they were not (Figure 6A), showing that the Arp2/3 complex is crucial for actin polymerization at damaged lysosomes.

**Figure 6.**
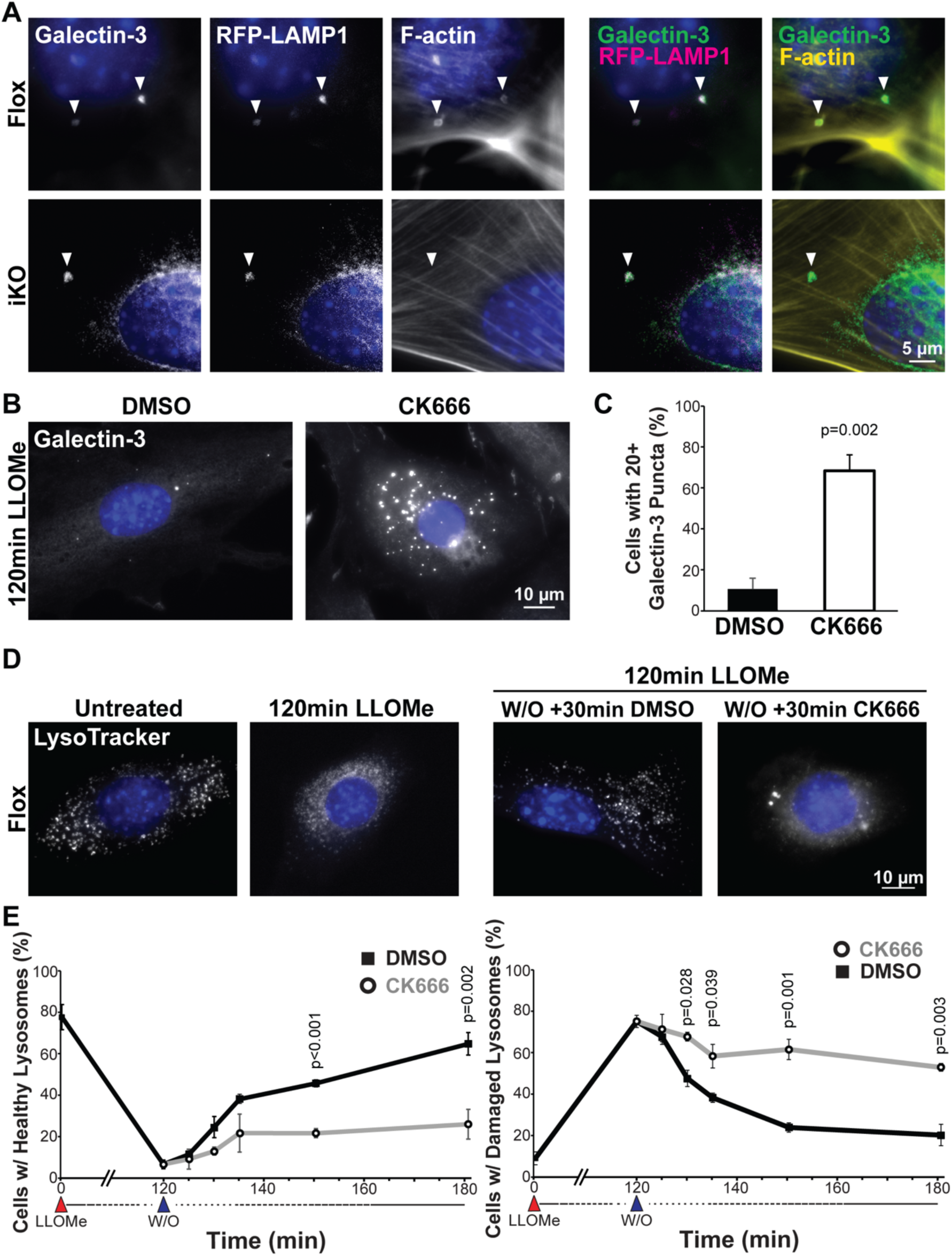
Arp2/3 complex-mediated actin assembly is important for lysosome repair. **(A)** MTFs expressing RFP-LAMP1 (magenta) were treated with DMSO (Flox) or 4-OHT (iKO) for 6d, exposed to 1mM LLOMe for 15 min, fixed, and stained with antibodies to detect Galectin-3, with phalloidin to visualize filamentous-actin (F-actin), and with DAPI. **(B)** MTFs were exposed to LLOMe for 2h in media containing either DMSO or CK666, fixed, and stained with Galectin-3 antibodies and DAPI. **(C)** The percentage of cells with 20 or more Galectin-3 puncta was quantified. Each bar represents the mean ±SD from n = 3 experiments. **(D)** MTFs were exposed to LLOMe for 2h on day 6, subjected to a washout (W/O) with media containing either DMSO or CK666 for the indicated timepoints and LysoTracker for 30min, fixed, and imaged. **(E)** The % of cells with either a damaged/diffuse (<3 puncta) or healthy/punctate (>10 puncta) LysoTracker staining pattern was quantified. Each symbol represents the mean % of cells with the depicted LysoTracker phenotype ±SD from n = 3 experiments.

The rapid assembly of actin suggested that the Arp2/3 complex may help maintain lysosomal integrity. To assess the effects of transient Arp2/3 complex inhibition during acute lysosomal damage, we treated Flox cells with LLOMe for two hours and included either DMSO or the Arp2/3 complex inhibitor CK666 in the media prior to staining for Galectin-3 to observe damaged lysosomes. The DMSO-treated control cells contained several Galectin-3 puncta as expected, while the CK666-treated cells displayed signs of much more severe damage (Figure 6B). Approximately 10% of control cells contained 20 or more Galectin-3 puncta compared to nearly 70% of the cells exposed to CK666 (Figure 6C), indicating that inhibition of the Arp2/3 complex greatly increases the susceptibility of cells to LLOMe.

The increased sensitivity to acute lysosomal membrane damage suggested that the Arp2/3 complex might be important for lysosomal repair. We therefore performed lysosome repair assays in which LysoTracker was used for monitoring healthy lysosomes first at steady state, then following an extended LLOMe treatment (for two hours), and eventually after LLOMe washout for 5-60 minutes (Eriksson et al., 2020). To determine the contributions of the Arp2/3 complex specifically during the lysosomal repair period, we again used Flox cells and included DMSO or CK666 in the washout media. Before exposure to LLOMe, most cells had normal LysoTracker-stained lysosomes (Figure 6D), as 80% were scored as ‘healthy’, defined as having >10 LysoTracker puncta per cell, whereas only about 10% were categorized as ‘damaged’ with diffuse LysoTracker staining, defined as having <3 puncta per cell (Figure 6E).

As expected, following LLOMe treatment, the scoring pattern was reversed, because nearly 80% of cells were classified as damaged and <10% as healthy (Figure 6E). After the washout of LLOMe, cells incubated in DMSO-containing media demonstrated a gradual return to a normal lysosomal population, with 60% of cells scoring as healthy by 60 minutes (Figure 6E). In contrast, cells incubated in CK666-containing media were significantly slower in their recovery and did not regain their pre-LLOMe staining patterns, as <25% of cells were scored as healthy through the entire washout timecourse (Figure 6E). The increase in healthy lysosomes was not due to clearance of damaged lysosomes by lysophagy, as Galectin-3 staining demonstrated that the number of Galectin-3 puncta did not change during the 60 minute washout (Figure S2). These data reveal that the Arp2/3 complex is necessary for repair of damaged lysosomes.

The Arp2/3 complex is an inefficient actin nucleator by itself and must be activated by nucleation-promoting factors from the WASP family to effectively assemble branched actin networks (Alekhina et al., 2017; Campellone and Welch, 2010). To better understand the mechanism underlying Arp2/3 activation in the lysosome repair response, we employed a panel of knockout (KO) human fibroblast-like cell lines that were engineered to separately lack each nucleation-promoting factor – N-WASP, WAVE1, WAVE2, WAVE3, the WAVE Complex, the WASH Complex, WHAMM, JMY, both WHAMM and JMY, Cortactin, or WISH/DIP1/SPIN90 (King et al., 2021). Two parental cell lines – nearly-haploid HAP1 cells and fully-haploid eHAP cells – served as controls, as did a KO cell line engineered to lack the non-cytoskeletal protein WDR73 (Essletzbichler et al., 2014; Mathiowetz et al., 2017). After confirming that F-actin could be assembled at damaged lysosomes in HAP1 and eHAP cells (Figure S3), we subjected the parental and KO cells to an extended LLOMe treatment (for three hours), stained them with Galectin-3 antibodies, and examined them by fluorescence microscopy (Figure 7A; Figure S3). Among the 14 cell lines that were analyzed, three knockouts showed significant changes in punctate Galectin-3 staining compared to their respective parental cell lines. WASH complex KO cells were devoid of puncta, whereas WHAMM KO cells had slightly more puncta, and WHAMM/JMY double knockout (DKO) cells had the greatest increase in puncta quantities (Figure 7A and 7B). Such differences in Galectin-3 accumulation at lysosomes led us to explore potential roles for WASH, WHAMM, and JMY in the lysosome damage response.

**Figure 7.**
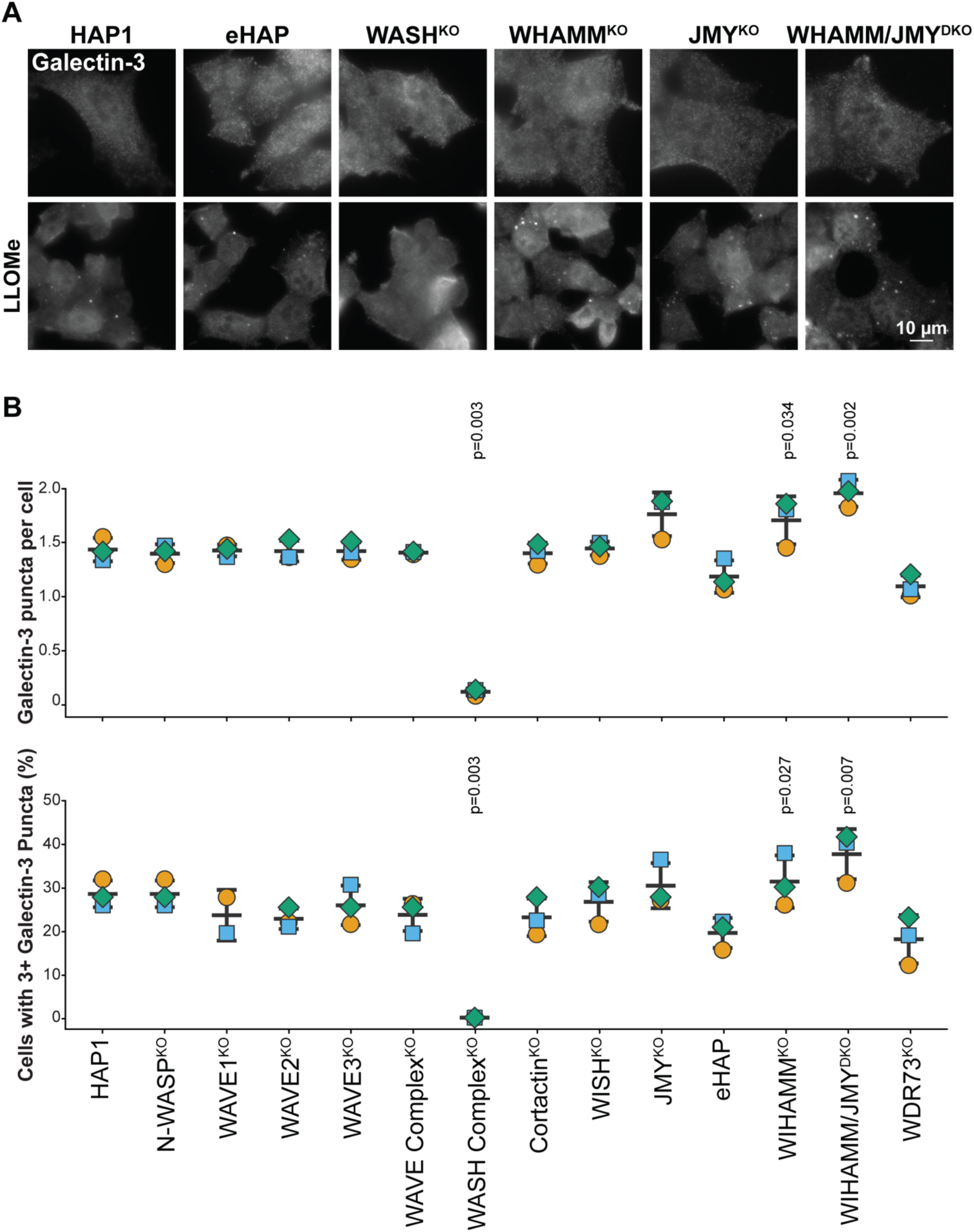
WHAMM/JMY double knockout (DKO) cells experience increased lysosomal damage following LLOMe treatment. **(A)** Parental (HAP1 or eHAP) or CRISPR-engineered knockout (KO) human fibroblasts lacking one or more Arp2/3-activating proteins were cultured in either normal growth media or media containing 0.5mM LLOMe for 3h before being fixed and stained with antibodies to Galectin-3. **(B)** The numbers of Galectin-3 puncta per cell were quantified. Symbols represent the average #s from individual experiments in which 57-109 cells were examined. Lines denote the mean ±SD from n = 3 experiments.

These three nucleation factors have distinct intracellular localizations. WASH is known to localize to endosomes and lysosomes (Derivery et al., 2009; Duleh and Welch, 2010; Gomez and Billadeau, 2009). WHAMM can be found at the ER, ER-Golgi intermediate compartment (ERGIC), and *cis*-Golgi at steady state, or at autophagosomes during nutrient starvation (Campellone et al., 2008; Kast et al., 2015; Mathiowetz et al., 2017; Russo et al., 2016). JMY is predominantly nuclear, but can be recruited to the plasma membrane, *trans*-Golgi, autophagosomes, or intrinsic apoptotic signaling structures depending on the cell type and experimental stressor (Coutts and La Thangue, 2015; King and Campellone, 2023; King et al., 2021; Schlüter et al., 2014; Shikama et al., 1999; Zuchero et al., 2009). To define the localization of these nucleation factors under normal and lysosome-damaging conditions, we stained HAP1 and eHAP cells with antibodies to LAMP2 and either WASH, WHAMM, or JMY.

As expected, in normal culture settings, WASH localized to cytoplasmic vesicles and a subset of LAMP2-positive lysosomes, JMY was found primarily in the nucleus, and WHAMM displayed a cytoplasmic and perinuclear localization pattern including occasional small puncta (Figure 8A). Upon exposure of cells to LLOMe, WASH localization to lysosomes decreased, JMY remained nuclear, and WHAMM underwent a striking redistribution onto the surface of nearly every lysosome in every cell (Figure 8A and 8B). More than 70% of WASH puncta were associated with LAMP2-positive lysosomes at steady state compared to <30% after LLOMe, whereas WHAMM recognition at lysosomes went from <5% in control conditions to nearly 95% after LLOMe-induced damage (Figure 8C). These observations highlight the different intracellular dynamics of WASP-family proteins and support the conclusion that WHAMM is the Arp2/3 complex activator directly involved in the cytoskeletal response to lysosomal damage.

**Figure 8.**
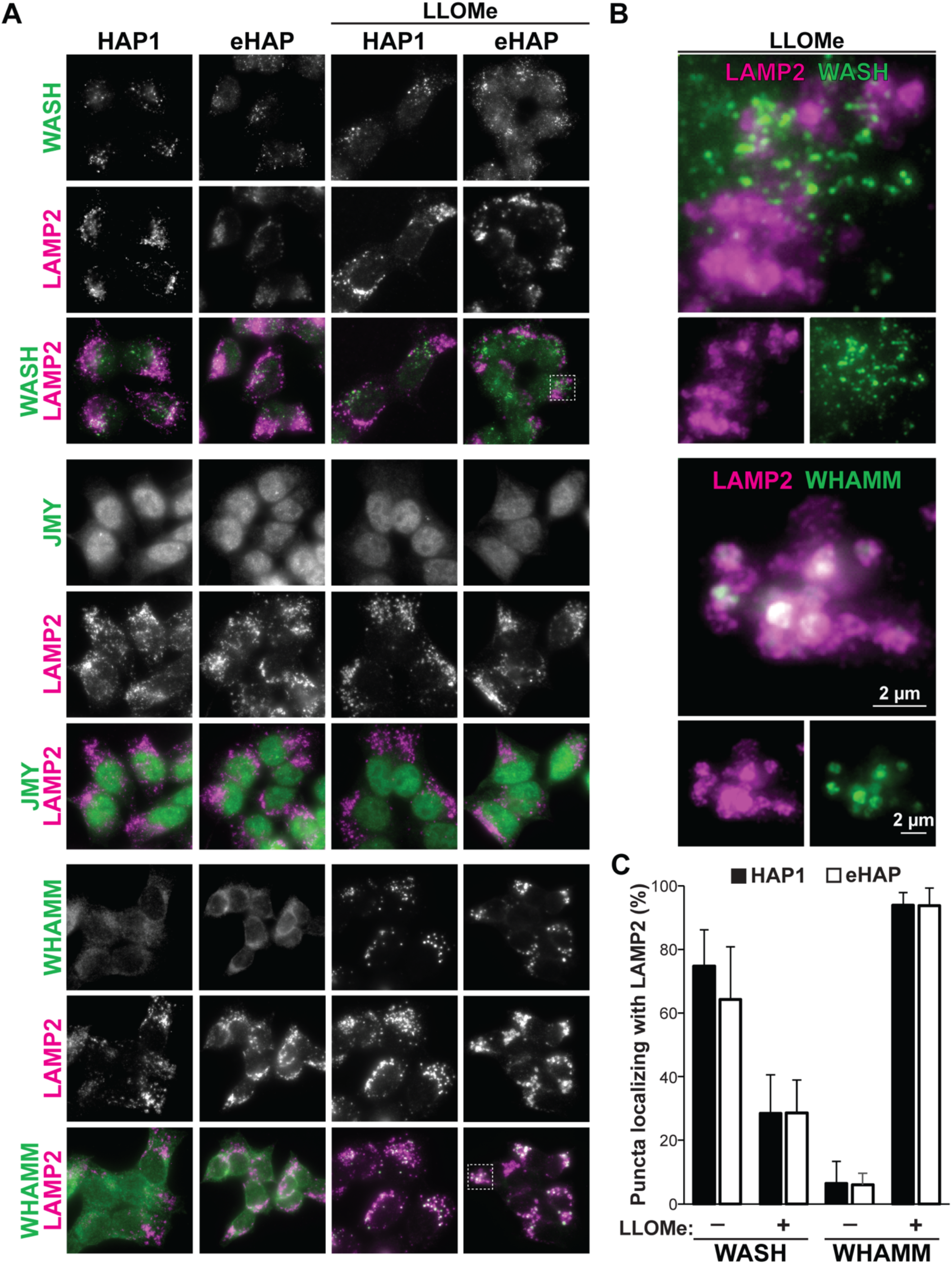
WHAMM is recruited to lysosomes following LLOMe-induced damage. **(A)** HAP1 and eHAP cells were cultured in normal media or treated with 0.5mM LLOMe for 3h, fixed, and stained with antibodies to LAMP2 (magenta) and either WASH, JMY, or WHAMM (green). **(B)** Magnified images of boxes highlighted in (A). **(C)** The percentage of WASH or WHAMM puncta that co-stained for LAMP2 was quantified. Bars represent the average %s ±SD from independent experiments in which n = 53-56 cells were examined.

## DISCUSSION

The Arp2/3 complex is most often recognized for its central role in actin assembly at the plasma membrane, but emerging evidence has drawn attention to its involvement in the process of autophagy. Several WASP-family proteins promote Arp2/3-mediated actin assembly during numerous aspects of the canonical autophagy pathway, from phagophore biogenesis to autophagosome motility to autolysosome tubulation, but the overall consequence of Arp2/3 inactivation has remained unclear. Using inducible ArpC2 knockout mouse fibroblasts, we now show that Arp2/3 complex activity is most important at the lysosome and that WHAMM is the most prominent WASP-family member which regulates Arp2/3-mediated actin assembly during a lysosomal damage response. Our results give rise to a model in which the Arp2/3 complex can polymerize actin to help maintain lysosomal integrity during basal growth conditions, and that stresses which induce more severe lysosomal damage trigger WHAMM-driven Arp2/3 activation to enable membrane repair (Figure 9).

**Figure 9.**
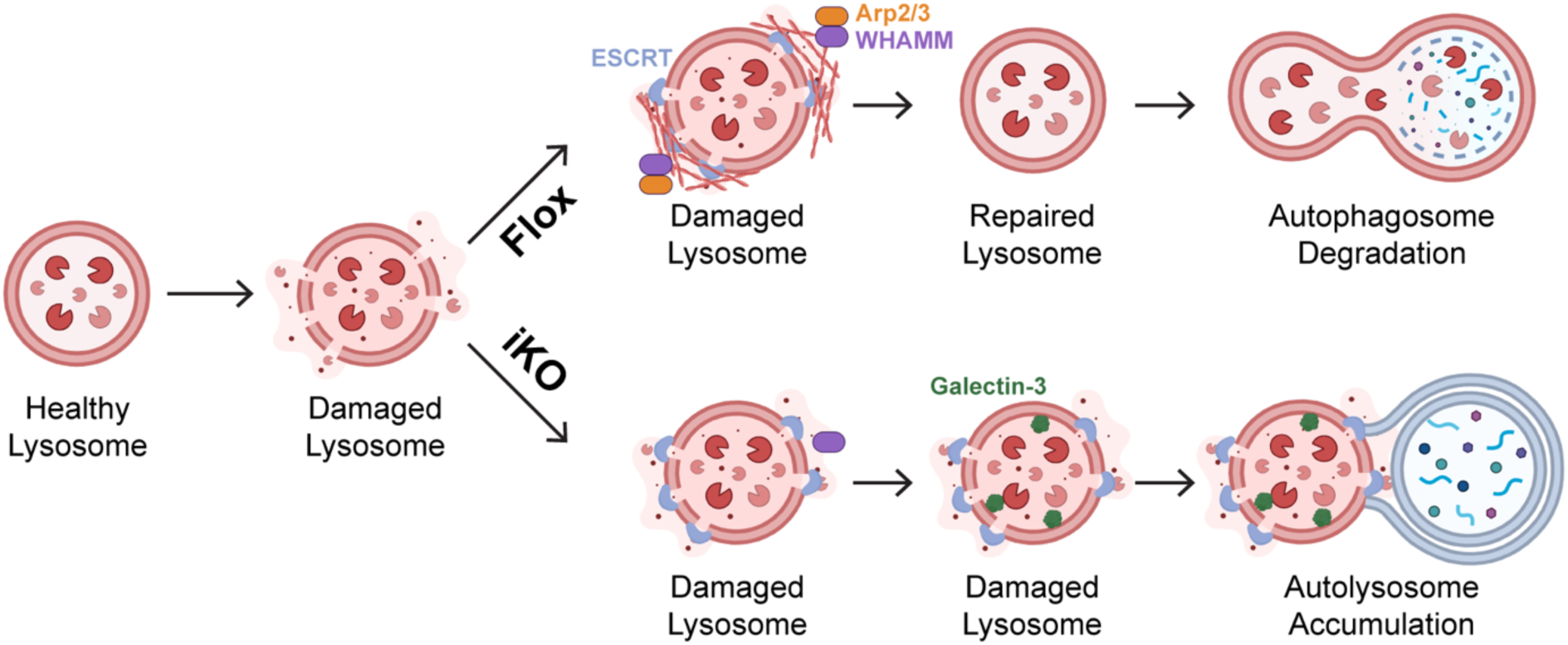
Model for Arp2/3 complex-mediated lysosomal maintenance in autophagy. Upon lysosomal damage, WHAMM localizes to the lysosome in normal cells (Flox) and activates the Arp2/3 complex to nucleate actin filaments which mediate lysosomal repair, allowing degradation of autophagosomes during autophagy. In cells lacking the Arp2/3 complex (iKO), actin is not assembled at the lysosome, leading to an increase in recruitment of Galectin-3 and an accumulation of damaged lysosomes which are unable to degrade autophagosomes.

A variety of localization and loss-of-function studies indicate that the Arp2/3 complex can be activated by WHAMM, JMY, WASH, Cortactin, and perhaps other nucleation-promoting factors during autophagy (Campellone et al., 2023; Kramer et al., 2022). However, differences in cell lines, fluorescent probes, overexpression levels, knockdown efficiencies, and unknown compensatory changes in knockout cells have made it difficult to elucidate when and where Arp2/3 complex activity is truly needed. By utilizing the ArpC2 iKO mouse fibroblast system (Rotty et al., 2015), we avoided complications from incomplete protein depletion and found that the net outcome of Arp2/3 complex ablation is a defect in autolysosome turnover. Analyses of both the LC3 and GABARAP subclasses of the ATG8 family, as well as fluorescent reporters of autophagosome and lysosome acidification, support the conclusion that mature autophagosomes are unable to be properly degraded by lysosomes in Arp2/3-deficient cells.

Careful quantification revealed that this dominant phenotype overshadows an inefficiency in autophagosome biogenesis. While our work emphasizes that Arp2/3 complex-dependent actin assembly is not equally important at each step of the autophagic pathway in fibroblasts, it does not detract from previous findings that Arp2/3 activators from the WASP-family are more specifically distributed. On the contrary, given that the rates of autophagic organelle biogenesis and turnover are different in distinct cell types (David et al., 2021; Evans and Holzbaur, 2020), it highlights the need for future studies to define which WASP-family members are the most relevant in specialized cells and physiological systems.

Aside from clarifying the importance of Arp2/3 complex function in autophagy, our work uncovers a new role for this actin nucleator in lysosomal repair. The presence of an unhealthy population of lysosomes in the absence of the Arp2/3 complex was first suggested by the observation of diffuse LysoTracker staining in senescent iKO cells (Haarer et al., 2023). In our current work, we confirmed the change in LysoTracker distribution in iKO cells at an earlier timepoint and additionally found that IST1, a member of the ESCRT-III repair complex (Radulovic et al., 2018), and Galectin-3, a lectin that recognizes the inside of damaged lysosomal membranes (Jia et al., 2020), were both more frequently associated with lysosomes when the Arp2/3 complex was deleted. Moreover, when healthy cells were treated with LLOMe to induce severe lysosomal damage, the simultaneous pharmacological inhibition of Arp2/3 with CK666 caused a major spike in recruitment of Galectin-3 to lysosomes. The finding that CK666 prevented the re-acidification of discrete lysosomes after LLOMe washout parallels the earlier result that lysosomes could not quench GFP-LC3 after chloroquine washout.

While we do not exclude the possibility that the Arp2/3 complex also has some impact on autophagosome-lysosome fusion, we conclude that the Arp2/3 complex is crucial for maintaining lysosomal integrity and is capable of responding to severe membrane damage by enabling lysosomal membrane repair pathways. This idea is again supported by LLOMe washout experiments wherein CK666 prevented the restoration of intact LysoTracker-stained organelles. We considered that the Arp2/3 complex might be affecting lysophagy in our cellular system, but Galectin-3 puncta were not cleared in the 10-minute to 3-hour range of recovery times that we examined, and lysophagy in other cell lines typically takes 6-24 hours (Gallagher and Holzbaur, 2023; Maejima et al., 2013). As implied by another study (Kravić et al., 2022), it remains likely that the Arp2/3 complex participates in lysophagy, but such functions for the actin nucleation machinery would occur beyond the times assayed in the above experiments. Our observations of Arp2/3 complex activity early in a lysosomal damage response, when taken together with other pharmacological studies of Arp2/3 during autophagic lysosome reformation (Dai et al., 2019; Wu et al., 2021) and lysophagy (Kravić et al., 2022), indicate that Arp2/3 complex deficiency creates a vicious cycle that leaves the cellular lysosome population in a largely unrepairable, unreformable, and undegradable state.

Although the roles of actin assembly in lysosome reformation and lysophagy still require further investigation, we focused on the mechanism of Arp2/3 activation during the short-term responses to acute lysosomal damage. Using a panel of human knockout fibroblasts lacking every characterized Arp2/3 complex activator (King et al., 2021), we found that WHAMM, WASH, and perhaps to a lesser extent, JMY, influence the Galectin-3 response to LLOMe-damaged lysosomes. Under the conditions we tested, JMY did not localize to lysosomes, so we speculate that JMY may be transiently recruited to damaged lysosomes in specialized circumstances to modulate a repair response and/or affect a transcriptional program from its position in the nucleus to increase lysosome biogenesis *de novo*. Consistent with early studies (Derivery et al., 2009; Duleh and Welch, 2010; Gomez et al., 2012), we found that WASH could localize to some lysosomes under steady state culture conditions. Surprisingly, however, the association of WASH with lysosomes decreased following LLOMe-induced damage, and WASH complex-depleted cells were mostly devoid of punctate Galectin-3 staining. WASH is known to influence the size, shape, position, and function of lysosomes (Gomez et al., 2012; King et al., 2013; Park et al., 2013; Priya et al., 2023). Given such previously-described roles for WASH in endolysosomal membrane protein trafficking and in the delivery of digestive hydrolases to lysosomes, the absence of punctate Galectin-3 staining in WASH complex-deficient cells could be reasonably explained by a failure of LLOMe to be processed by cathepsins and a lack of lumen-facing glycoproteins that could be recognized by Galectin-3. Hence, we were more intrigued by the striking accumulation of WHAMM at lysosomes upon LLOMe-induced damage.

WHAMM participates in many aspects of intracellular trafficking, from ER-Golgi transport and membrane tubulation (Campellone et al., 2008; Russo et al., 2016), to autophagosomal membrane remodeling (Coulter et al., 2024; Kast et al., 2015; Mathiowetz et al., 2017), to post-mitochondrial apoptotic signaling (King and Campellone, 2023; King et al., 2021). The highly penetrant localization of WHAMM to lysosomes specifically after LLOMe exposure supports the model that WHAMM is the key nucleation-promoting factor that responds to membrane damage and mediates Arp2/3 complex activation. The additional findings that deletion of human WHAMM, either by itself or in combination with JMY, results in increased Galectin-3 staining upon LLOMe-driven damage, suggest that like Arp2/3, WHAMM is functionally necessary for managing the repair of severely damaged lysosomal membranes. The reasons why mouse Arp2/3 iKO fibroblasts harbored greater steady-state levels of Galectin-3 puncta compared to human WHAMM KO or WHAMM/JMY DKO fibroblasts are unclear, but could be due to species- or tissue-specific differences, or from compensatory gene expression changes that accompany long-term nucleation factor inactivation. It is also possible that the Arp2/3 complex is regulated by other proteins in addition to WHAMM and JMY to mitigate multiple intracellular stresses. The extent to which WHAMM controls Arp2/3 localization and lysosomal damage in mouse cells has yet to be established. The broader mechanisms that govern the spatiotemporal dynamics of nucleation factors also warrant future investigation, including how WASH is displaced from lysosomes, how JMY impacts lysosomal function, how WHAMM is recruited to lysosomes, and how the Arp2/3 complex affects specific membrane repair activities.

Recent years have produced many advances in understanding lysosomal repair and reformation mechanisms. These include a damage recovery process mediated by ESCRT complexes (Radulovic et al., 2018; Skowyra et al., 2018), a repair pathway involving lipid transport at ER contact sites (Radulovic et al., 2022; Tan and Finkel, 2022), and a regeneration method associated with ATG8 (Bhattacharya et al., 2023). Determining where WHAMM, the Arp2/3 complex, and actin fit in the context of these or other lysosomal homeostasis mechanisms is an important avenue for future investigation. It will be interesting to see whether the cytoskeletal proteins play active roles in damage surveillance, organelle sequestration, structural maintenance, membrane shaping, scaffolding, or signaling.

Finally, it is important to note that actin assembly is emerging as a common feature of many cellular stress responses (Campellone et al., 2023). Our work now adds lysosomal stress to the genotoxic, nutritional, and proteostatic stresses that have already been shown to trigger actin reorganization. Perhaps comparisons will be drawn to the cytoskeletal rearrangements that influence mitochondrial homeostasis during normal quality control and in response to mitochondrial dysfunction (Chakrabarti et al., 2022; Fung et al., 2019; Moore and Holzbaur, 2021; Moore et al., 2016). At the organismal level, lysosomal damage responses are relevant to many diseases, including infections (Boyle and Randow, 2013), neurodegenerative disorders (Flavin et al., 2017), and other aging-associated conditions (Gómez-Sintes et al., 2016; Kirkegaard et al., 2010). Enhancing our understanding of the connections between the actin nucleation machinery and lysosomal function may therefore provide opportunities for developing molecular interventions that manipulate organelle repair mechanisms during aging and disease.

## MATERIALS AND METHODS

### Mammalian cell culture

All cell lines are listed in Table S1. Mouse tail fibroblasts (MTFs) containing the floxed *Arpc2* allele (from James Bear, University of North Carolina) (Rotty et al., 2015) were cultured in DMEM (with 4.5g/L glucose, L-Glutamine, 110mg/L sodium pyruvate), 10% fetal bovine serum (FBS), GlutaMAX, and antibiotic-antimycotic (Gibco). Media containing 0.01% DMSO or 2µM 4-OHT (Sigma) was used to generate Flox or iKO populations. For treatments exceeding 3 days, cultures were replenished with fresh media containing DMSO or 4-OHT on day 4 and returned to normal media after day 6. HAP1 cell derivatives were cultured in Iscove’s Modified Dulbecco’s Medium (IMDM; Invitrogen), 10% FBS, and penicillin-streptomycin, as described previously (King et al., 2021). All cell lines were grown at 37°C in 5% CO_2_, and assays were performed using cells that had been in active culture for 2-10 trypsinized passages.

### DNA transfections and fluorescent probes

For transient tfLC3 expression, MTFs were cultured in 6-well plates for 24h before media with DMSO or 4-OHT was added, and cells were treated for 48h before transfection. Both Flox and iKO cells were transfected with 600ng of tfLC3 plasmid (Addgene, 21704) using Lipofectamine LTX (Invitrogen) in DMEM. After 4h, DMEM was replaced with MTF media, and 18h later the cells were trypsinized and transferred onto 12mm glass coverslips in 24-well plates. For stable LAMP1-mRFP expression, a LAMP1-mRFP plasmid (Addgene, 34611) was linearized using NdeI, and 500ng was transfected into MTFs. After 4h, DMEM was replaced with MTF media, and 18h later the cells were trypsinized and transferred into 10cm dishes with media containing 1μg/mL puromycin. Cells were grown for 6d before individual colonies were trypsinized and transferred to 12-well plates. To identify fluorescent clones, cells were fixed onto glass coverslips and viewed microscopically. One representative clone was used in experiments. For imaging acidic cytoplasmic organelles, cells were incubated for 30min in media containing 50nM LysoTracker Red (Invitrogen) prior to fixation.

### Chemical treatments

For starvation experiments, Flox and iKO cells were grown on glass coverslips in 24-well plates. Normal MTF culture media was replaced with Hanks’ Balanced Salt Solution (Gibco) for 3.5h before the cells were fixed and stained as described below. For mitochondrial depolarization, MTF media was replaced with media containing 10μM CCCP (Tocris) for 3h. For lysosome inhibition, normal growth media was replaced with media containing 50μM chloroquine (Sigma) for 2h. For autophagosome turnover assays, the chloroquine media was removed and replaced with normal media for 15min to 4h. For lysosome damaging experiments, cells were treated with 0.5 or 1.0mM LLOME (MedChemExpress) for 15min to 3h. LLOMe undergoes cathepsin C-dependent processing and polymerization to generate its membranolytic species (Kavčič et al., 2020; Thiele and Lipsky, 1990). For lysosome recovery assays, cells were treated with media containing 1mM LLOMe for 2h, then the LLOMe media was replaced with normal media containing either 50μM CK666 (Calbiochem) or an equivalent volume of DMSO for 0-60min before fixation. For the 0-15min recovery timepoints, LysoTracker was added to the LLOMe-containing media 30min prior to fixation. For the 30-60min timepoints, LysoTracker was added to the DMSO or CK666-containing media 30min prior to fixation.

### Immunoblotting

For preparation of cell extracts, MTFs cultured in 6-well plates were collected in phosphate-buffered saline (PBS) containing 1mM EDTA, centrifuged at 750 x g for 5.5min at 4°C, and lysed in 25mM HEPES (pH 7.4), 100mM NaCl, 1% Triton-X-100, 1mM EDTA, 1mM Na_3_VO_4_, 1mM NaF, 1mM PMSF, and 10μg/ml each of aprotinin, leupeptin, pepstatin, and chymostatin on ice. Lysates were mixed with Laemmli sample buffer, boiled, and centrifuged. Extract samples were separated on 12% SDS-PAGE gels before transfer to nitrocellulose membranes (GE Healthcare). Membranes were blocked in PBS containing 5% milk before probing with primary antibodies at concentrations listed in Table S2. Following overnight incubation at 4°C, membranes were washed and treated with IRDye-680/800-(LI-COR) or horseradish peroxidase-conjugated (GE Healthcare) secondary antibodies. Infrared and chemiluminescent bands were visualized using a LI-COR Odyssey Fc Imaging System. Band intensities were measured using LI-COR Image Studio software. Densitometries of proteins-of-interest were normalized to GAPDH, tubulin, and/or actin loading controls.

### Immunostaining

For immunofluorescence, fibroblasts cultured on glass coverslips in 24-well plates were washed with PBS and fixed using PBS containing 2.5% paraformaldehyde for 20-35min. Following PBS washes, cells were permeabilized using PBS containing 0.1% Triton X-100 for 2min, washed, and stained with primary antibodies in PBS containing 1% bovine serum albumin, 1% FBS, and 0.02% azide for 45-60min as described in Table S2. Cells were washed and treated with Alexa Fluor 488-, 555-, or 647-conjugated goat anti-rabbit or anti-mouse secondary antibodies, 1μg/mL DAPI, and/or 0.2U/mL Alexa Fluor 647-labeled phalloidin (Invitrogen) for 35-45min as detailed in Table S2. Following washes, coverslips were mounted on glass slides in ProLong Gold anti-fade (Invitrogen).

### Fluorescence microscopy

All cells were imaged using a Nikon Eclipse Ti microscope equipped with Plan Apo 100X (1.45 NA), Plan Apo 60X (1.40 NA), or Plan Fluor 20X (0.5 NA) objectives, an Andor Clara-E camera, and a computer running NIS Elements Software. All cells were viewed in multiple focal planes, and Z-series were captured at 0.2–0.4μm steps. Images presented in the figures represent one slice (Figs 2,6,7) or multiple-slice maximum intensity projections (Figs 3,4,5,8).

### Image processing and quantification

All image processing and analysis was conducted using ImageJ/FIJI software (Schindelin et al., 2012). The ImageJ Cell Counter plug-in was used to quantify the percentage of cells with LC3 or GABARAP-associated autophagosomes by manually counting the total cells in the DAPI channel and the cells that were positive for cytosolic rings in the LC3 or GABARAP channels.

For the number of autophagosomes per cell, the ImageJ Dot Counter tool was used to manually mark the individual LC3 or GABARAP rings in each cell. The same method was used for the number of IST1 and Galectin-3 puncta per cell. For the number of tfLC3 autophagosomes and autolysosomes, the Cell Counter plug-in was used to mark the white autophagosomes versus magenta autolysosomes.

### Reproducibility and statistics

All conclusions were based on observations made from at least 3 separate experiments, while quantifications were based on data from 2-5 representative experiments. Statistical analyses were performed using SuperPlotsOfData software (Goedhart, 2021). P-values for comparisons of pairwise data sets were determined using paired t-tests while other data sets were analyzed using unpaired t-tests (Zweifach, 2024). P-values <0.05 were considered statistically significant.

## ACKNOWLEDGEMENTS

We thank Jim Bear (University of North Carolina) for providing cells harboring the floxed *Arpc2* allele and Campellone Lab members for their comments on this manuscript. KGC was supported by National Institutes of Health grants GM107441 and AG050774 (www.nih.gov). The funders had no role in study design, data collection and analysis, decision to publish, or preparation of the manuscript.

