## Supplemental Information for "Autophagosome turnover requires Arp2/3 complex-mediated maintenance of lysosomal integrity"

**SUPPLEMENTARY MATERIALS**

| **Table S1. Cell Lines** | |
|---|---|
| ***Parental Cells*** | |
| Cell Line | Source |
| eHAP | King *et al.,* 2021 |
| HAP1 | King *et al.,* 2021 |
| ArpC2 floxed MTFs | Rotty *et al.,* 2015 |
| ***HAP1 Derivatives*** | |
| KO Cell Line | Source |
| Cortactin$^{KO}$ | King *et al.,* 2021 |
| JMY$^{KO}$ | King *et al.,* 2021 |
| N-WASP$^{KO}$ | King *et al.,* 2021 |
| WASH Complex$^{KO}$ (CCDC53$^{KO}$) | King *et al.,* 2021 |
| WAVE1$^{KO}$ | King *et al.,* 2021 |
| WAVE2$^{KO}$ | King *et al.,* 2021 |
| WAVE3$^{KO}$ | King *et al.,* 2021 |
| WAVE Complex$^{KO}$ (BRK1$^{KO}$) | King *et al.,* 2021 |
| WISH$^{KO}$ | This study;<br>8bp deletion in exon 2 of *WISH/DIP1/SPIN90*,<br>Horizon Genomics (HZGHC003331c004) |
| ***eHAP Derivatives*** | |
| KO Cell Line | Source |
| WDR73$^{KO}$ | Mathiowetz *et al*., 2017 |
| WHAMM$^{KO}$ | Mathiowetz *et al*., 2017 |
| WHAMM/JMY$^{DKO}$ | King *et al*., 2021 |

**Table S2.** **Immunofluorescence and Immunoblotting Reagents**

| Target | Probe | | Conc. | Source |
|---|---|---|---|---|
| ***Primary Antibodies (Immunofluorescence)*** | | | | |
| GABARAP | anti-GABARAP | Rabbit | 1:750 | Proteintech (18723-1-AP) |
| Galectin-3 | anti-Galectin-3 | Rabbit | 1:1000 | Proteintech (14979-1-AP) |
| IST1 | anti-IST1 | Rabbit | 1:1000 | Proteintech (19842-1-AP) |
| JMY | anti-JMY | Rabbit | 1:1000 | Proteintech (25098-1-AP) |
| LAMP2 | anti-CD107b | Mouse | 1:500 | Proteintech (65053-1-Ig) |
| LC3 | anti-LC3 | Rabbit | 1:750 | Proteintech (14600-1-AP) |
| WASH | anti-WASH | Rabbit | 1:1000 | Duleh and Welch, 2010 |
| WHAMM | anti-WHAMM | Rabbit | 1:1000 | Shen *et al.*, 2012 |
| ***Primary Antibodies (Immunoblotting)*** | | | | |
| Actin | anti-Actin | Mouse | 1:10,000 | Proteintech (66009-1-Ig) |
| ArpC2 | anti-ArpC2 | Rabbit | 1:1000 | Millipore (07-227-I) |
| GABARAP | anti-GABARAP | Rabbit | 1:750 | Proteintech (18723-1-AP) |
| GAPDH | anti-GAPDH | Mouse | 1:10,000 | Proteintech (60004-1-Ig) |
| LC3 | anti-LC3 | Rabbit | 1:750 | Proteintech (14600-1-AP) |
| NBR1 | anti-NBR1 | Rabbit | 1:2000 | Proteintech (16004-1-AP) |
| NDP52 | anti-NDP52 | Rabbit | 1:2000 | Proteintech (12229-1-AP) |
| Optineurin | anti-Optineurin | Rabbit | 1:2000 | Proteintech (10837-1-AP) |
| SQSTM1/p62 | anti-SQSTM1/p62 | Rabbit | 1:1000 | Proteintech (18420-1-AP) |
| TAX1BP1 | anti-TAX1BP1 | Rabbit | 1:2000 | Proteintech (14424-1-AP) |
| Tubulin | anti-Tubulin | Mouse | 1:10,000 | DSHB (E7) |
| ***Secondary Antibodies (Immunofluorescence)*** | | | | |
| Mouse IgG | Alexa 488, 555, 647 anti-mouse | Goat | 4 μg/ml | Life Technologies (e.g.A11029) |
| Rabbit IgG | Alexa 488, 555, 647 anti-rabbit | Goat | 4 μg/ml | Life Technologies (e.g.A11034) |
| ***Secondary Antibodies (Immunoblotting)*** | | | | |
| Rabbit IgG | HRP anti-Rabbit | Donkey | 1:10,000 | GE Healthcare (NXA931) |
| Mouse IgG | IRDye 680, 800 anti-Mouse | Donkey | 0.05 μg/ml | LI-COR (e.g. 926-32212) |
| Rabbit IgG | IRDye 680, 800 anti-Rabbit | Donkey | 0.05 μg/ml | LI-COR (e.g. 926-32213) |
| ***Molecular Probes (Fluorescence)*** | | | | |
| Target | Probe | | Conc. | Source |
| DNA | 4’,6-diamidino-2-phenylindole (DAPI) | | 1 μg/ml | Invitrogen (D1306) |
| F-actin | Alexa 647-Phalloidin | | 0.2U/ml | Invitrogen (A22287) |
| Lysosomes | LysoTracker Red | | 50nM | Invitrogen (L7528) |

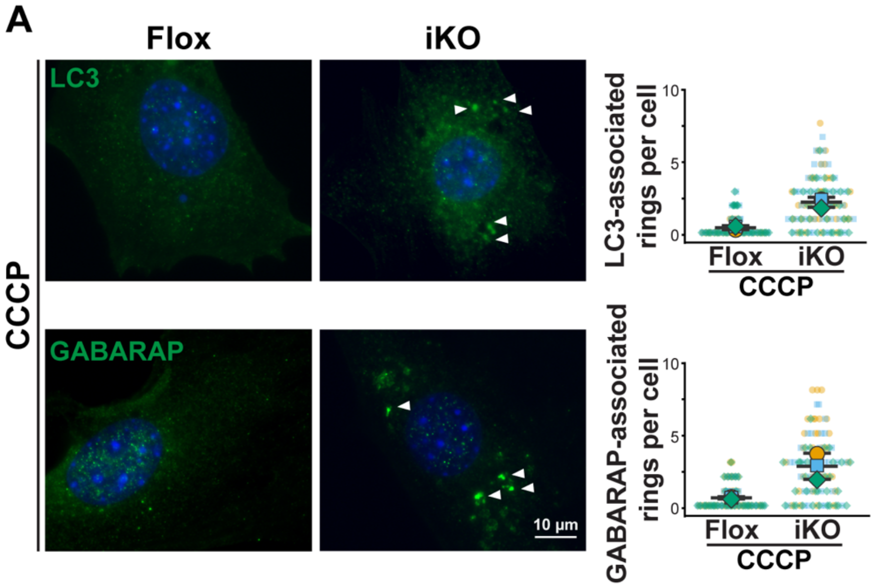


**Supplemental Figure S1. Deletion of the Arp2/3 complex leads to an increase in mature autophagic rings in cells with damaged mitochondria. (A)** MTFs were treated with DMSO (Flox) or 4-OHT (iKO) for 6d and exposed to 10μM CCCP for 3h on day 8. Cells were then fixed and stained with antibodies to detect LC3 or GABARAP (green) and with DAPI to visualize DNA (blue). Arrowheads highlight mature autophagosomal rings. **(B)** Superplots depict quantification of the number of mature autophagosomal rings per cell. Small symbols represent the #s within individual cells. Large symbols represent the average #s from individual experiments in which 32-38 cells were examined. Lines denote the mean ±SD from n = 3 experiments.

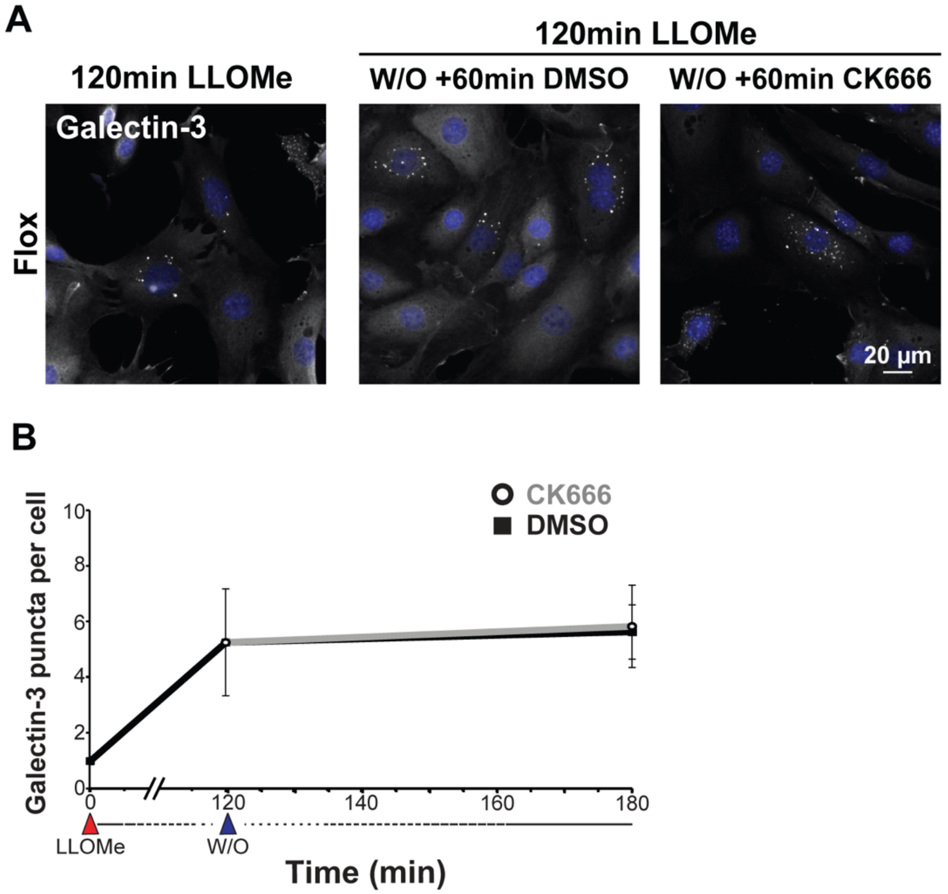


**Supplemental Figure S2. Arp2/3 complex inhibition does not influence the short-term clearance of lysosomes. (A)** Flox cells were exposed to 1mM LLOMe for 2h on day 6, subjected to a washout (W/O) with media containing either DMSO or CK666 for the indicated timepoints and LysoTracker for 30min, fixed, stained with Galectin-3 antibodies (gray) and DAPI (blue), and imaged. **(B)** The number of Galectin-3 puncta per cell was quantified. Each point represents the average #s from individual experiments in which 57-60 cells were examined. Lines denote the mean ±SD from n = 3 experiments.

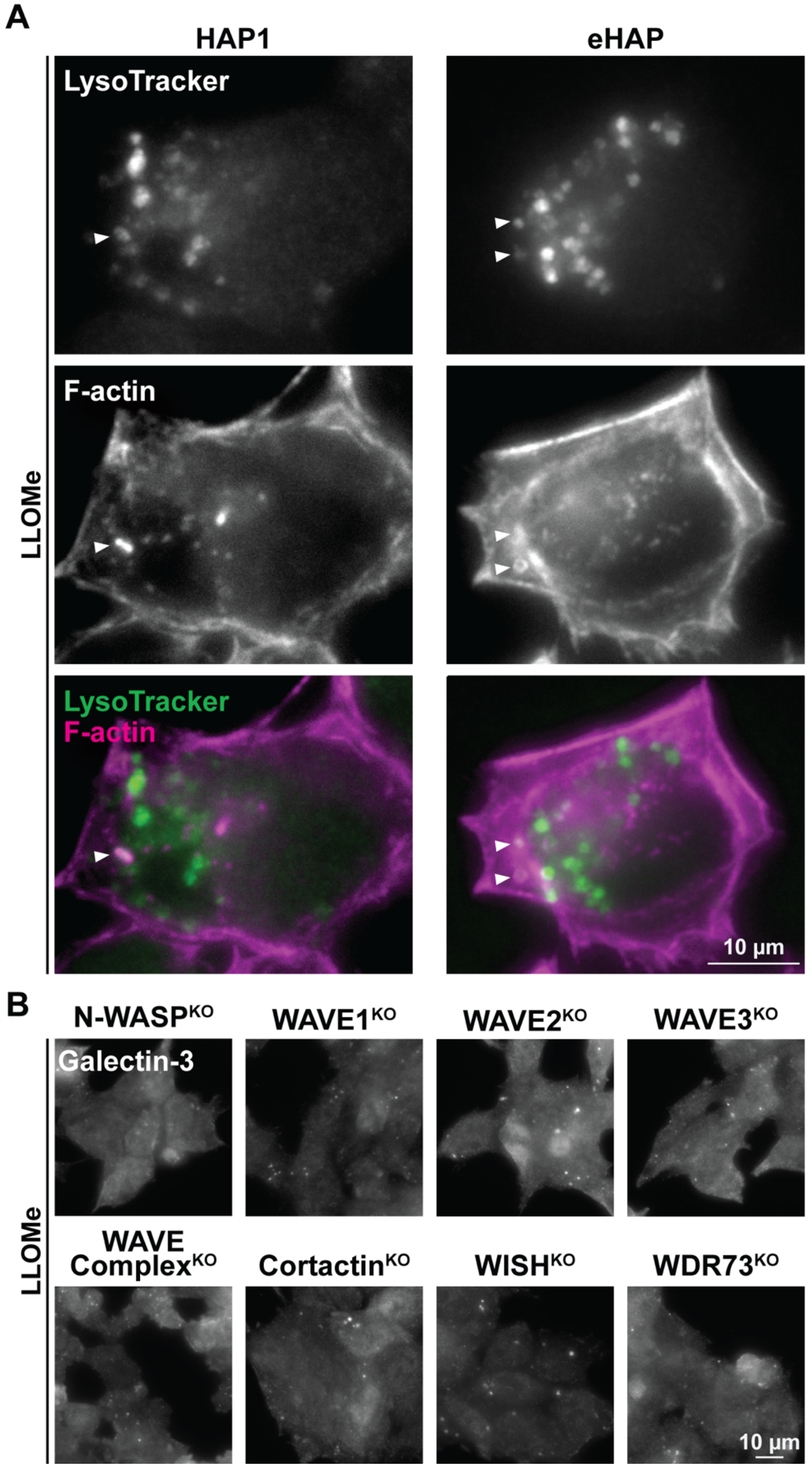


**Supplemental Figure S3. F-actin is recruited to damaged lysosomes in human fibroblasts and most Arp2/3-activating proteins do not influence the amount of lysosomal damage. (A)** HAP1 and eHAP human fibroblasts were treated with media containing 0.5mM LLOMe for 3h, with LysoTracker (pseudocolored green) for 30min, fixed, and stained with phalloidin to visualize F-actin (magenta). **(B)** CRISPR-engineered knockout human fibroblasts lacking Arp2/3-activating nucleation factors or a control protein (WDR73) were cultured in media containing LLOMe for 3h before being fixed and stained with antibodies to Galectin-3. These images accompany those presented in Figure 7.
